# The Metabolic Effects of Angiopoietin-like protein 8 (ANGPLT8) are Differentially Regulated by Insulin and Glucose in Adipose Tissue and Liver and are Controlled by AMPK Signaling

**DOI:** 10.1101/734954

**Authors:** Lu Zhang, Chris E. Shannon, Terry M. Bakewell, Muhammad A. Abdul-Ghani, Marcel Fourcaudot, Luke Norton

**Author notes:** Address all correspondence to: Luke Norton, PhD.

## Abstract

**Objective:** The angiopoietin-like protein (ANGPTL) family represents a promising therapeutic target for dyslipidemia, which is a feature of obesity and type 2 diabetes (T2DM). The aim of the present study was to determine the metabolic role of ANGPTL8 and to investigate its nutritional, hormonal and molecular regulation in key metabolic tissues.

**Methods:** The metabolism of ANGPTL8 knockout mice (ANGPTL8^−/−^) was examined in mice following chow and high-fat diets (HFD). The regulation of ANGPTL8 expression by insulin and glucose was quantified using a combination of *in vivo* insulin clamp experiments in mice and *in vitro* experiments in hepatocytes and adipocytes. The role of AMPK signaling was examined, and the transcriptional control of ANGPTL8 was determined using bioinformatic and luciferase reporter approaches.

**Results:** The ANGPTL8^−/−^mice had improved glucose tolerance and displayed reduced fed and fasted plasma triglycerides. However, there was no reduction in steatosis in ANGPTL8^−/−^mice after the HFD. Insulin acutely activated ANGPTL8 expression in liver and adipose tissue, which was mediated by C/EBPβ. Using insulin clamp experiments we observed that glucose further enhanced ANGPTL8 expression in the presence of insulin in adipocytes only. The activation of AMPK signaling potently suppressed the effect of insulin on ANGPTL8 expression in hepatocytes.

**Conclusion:** These data show that ANGPTL8 plays an important metabolic role in mice that may extend beyond triglyceride metabolism. The finding that insulin and glucose have distinct roles in regulating ANGPTL8 expression in liver and adipose tissue may provide important clues about the function of ANGPTL8 in these tissues.

## Introduction

Obesity and type 2 diabetes mellitus (T2DM) are associated with impaired lipid metabolism leading to elevated circulating free fatty acids (FFAs) and triglycerides. Dyslipidemia may play a causal role in the etiology of insulin resistance (1), and is a prominent risk factor for cardiovascular disease (2). Although current lipid-lowering medications can alleviate dyslipidemia, significant side effects associated with their use, including increased risk of T2DM and liver and muscle damage, highlight the need for new therapies that target lipid metabolism in metabolic diseases (3).

The angiopoietin-like protein (ANGPTL) family represents a promising therapeutic target for dyslipidemia. Two members of this family, ANGPTL3 and ANGPTL4, are well-established inhibitors of lipoprotein lipase (LPL), and loss of function of either protein leads to significant hypertriglyceridemia in mice (4). Furthermore, nonsense mutations in the human *ANGPTL3* gene cause familial combined hypolipidemia (FHBL2, OMIM #605019), characterized by extremely low levels of low-density lipoprotein (LDL) and high-density lipoprotein (HDL) cholesterol and triglycerides (5). Like other ANGPTL proteins, ANGPTL8 participates in the negatively regulation of LPL activity, and its inactivation similarly reduces serum triglyceride levels (6; 7). In human subjects a positive association between circulating ANGPTL8 and plasma triglycerides has been observed, and in genome-wide association studies (GWAS) polymorphisms in the *ANGPTL8* gene are associated with altered lipid profiles (8).

However, several important questions remain regarding the metabolic role of ANGPTL8 as well as its molecular regulation *in vitro* and *in vivo*. Firstly, whether ANGPTL8 influences adiposity and whole-body glucose homeostasis remains controversial. Initial studies in mice claimed ANGPTL8 regulated β-cell mass and function (9), but this finding was not supported in follow-up studies and was subsequently retracted (10). In mice lacking ANGPTL8, no changes in glucose tolerance were observed in two studies, but dramatic differences in the body weight and fat mass of ANGPTL8 knockout mice were reported in a third study (10). Similarly, in a more recent study ANGPTL8 inactivation using antisense oligonucleotides (ASO) increased fat mass and improved glucose tolerance in mice (11). In human studies circulating ANGPTL8 levels are elevated in patients with insulin resistance and T2DM, but this is also an inconsistent finding (12; 13). Secondly, the physiological and molecular mechanisms of ANGPTL8 gene regulation is relatively unexplored. This is an important area of research because the coordinated expression patterns of ANGPTL8 genes in response to nutritional status is likely a critical determinant of LPL activity and triglyceride trafficking, and may influence glucose metabolism. The expression of ANGPTL8 is highly abundant in the liver and adipose tissue, and is acutely regulated by nutritional status (14). Fasting and refeeding lead to a significant downregulation and upregulation of ANPTL8 expression, respectively (14), but the specific signaling mechanisms and the molecular pathways responsible in liver and adipose tissue remain to be elucidated.

In the following study we first examined the effect of ANGPTL8 knockout on glucose and lipid metabolism *in vivo*, and then investigated the physiological and molecular mechanisms regulating ANGPTL8 in adipose tissue and liver. Our metabolic studies in rodents confirm a powerful role for ANGPTL8 in regulating triglyceride metabolism but, in contrast to previous studies, demonstrate that ANGPTL8 knockout mice are partially protected from high-fat diet mediated glucose intolerance independent of any effects on bodyweight. We identify insulin and glucose as the key regulators of ANGPTL8 transcription, and describe important differences in the nutritional regulation of ANGPTL8 expression in adipose tissue and liver. Finally, we identify ANGPTL8 as a transcriptional target of CCAAT/enhancer-binding protein (C/EBPβ) in hepatocytes, and highlight the AMPK signaling pathway as a negative regulator of ANGPTL8 expression in the presence of inculsin. Taken together these studies reveal important new details about the regulation of ANGPTL8 *in vivo* and its effects on lipid and glucose metabolism.

## Materials and Methods

### ANGPTL8 knockout mice

Whole-body ANGPTL8 knockout mice (ANGPTL8^−/−^, B6; 129S5-Angptl8^tm1Lex^/Mmucd) were a gift from Dr. Ren Zhang at Wayne State University (Michigan, USA) and were originally obtained from the Mutant Mouse Resource Research Centers (7). Mice were fed either regular chow or a high-fat diet (HFD; Research Diets, D12492), where indicated. For fasting and refeeding experiments, mice were subjected to overnight (18 hours) food deprivation, followed by 6 hours refeeding. Littermate controls were used for all mouse experiments involving ANGPL8^−/−^ mice, and both male and female mice were examined.

### Mouse Physiological Studies

For intraperitoneal glucose tolerance tests (IPGTT) mice were fasted overnight for 12 hours and injected with glucose (2 g/kg) intraperitoneally. Blood glucose was measured on samples taken from the mouse tail at multiple time points for 120 minutes. Insulin clamp studies were carried out on wild-type C57BL/6J mice (stock number: 000664; Jackson Laboratory, Bar Harbor, ME, USA). Five days prior to clamp experiments, a catheter was inserted into the jugular vein of 12 – 14- week-old male mice. For euglycemic- and hyperglycemic-hyperinsulinemic clamps, the infusion of insulin (Humulin R) was initiated in a primed-continuous manner (50 mU/kg bolus followed by 5 mU/kg/min infusion). For hyperinsulinemic-somatostatin clamps, mice were infused with somatostatin at 3 μg/kg/min. Blood glucose was maintained at 100 mg/dL for euglycemic-hyperinsulinemic clamps, and at 300-350 mg/dL for both hyperglycemic clamps. To prevent volume depletion and anemia, red blood cells from a matched donor mouse were continuously infused during the clamp procedure.

### Insulin, Triglycerides and Free Fatty Acid Measurements

Mouse serum insulin levels were determined using the mouse metabolic hormone panel (Millipore Sigma, Milliplex MMHMAG-44K) according to the manufacturer’s instructions. Assays were performed on a MAGPIX Multiplexing instrument and data was analyzed using xPONENT® 4.2 software. Liver triglycerides were quantitated using the triglyceride quantification kit (Abcam, ab65336). Serum FFA levels were determined using a standard NEFA assay kit (Wako Chemicals USA, Inc., Richmond, VA, USA).

### Cell Culture studies

Both H4IIE and 3T3-L1 cells were obtained from the American Type Culture Collection (Rockville, MD, USA), and maintained in low glucose (5 mM) Dulbecco’s modified Eagle’s medium (DMEM). The 3T3-L1 cells were induced to differentiate into adipocytes before treatments, according to previous methodology (15). For studies of the regulation of ANGPTL8 gene expression by insulin and/or glucose, cells were cultured in serum-free DMEM supplemented with insulin and/or glucose for the indicated time points. All treatments in 3T3-L1 adipocytes started eight days after differentiation. For the insulin signaling inhibitor experiments, cells were pretreated for 60 minutes with vehicle (dimethyl sulphoxide, DMSO), 100 nM of the PI3K inhibitor wortmannin or 10 μM of the mitogen-activated protein kinase kinase (MAPKK) inhibitor PD98059, and were incubated for 2 hours in the presence of 100 nM insulin. For the AMPK activator 5-Aminoimidazole-4-carboxamide ribonucleotide (AICAR) treatment, cells were incubated with 0.5 mM AICAR in the presence of 100 nM insulin for two hours. Following treatments total mRNA and protein were extracted for qRT-PCR (**Table 1**) and immunoblotting assays (**Table 2**). All chemicals used in the *in vitro* experiments were obtained from Sigma-Aldrich (St. Louis, MO, USA).

**Table 1.** Real-time qRT-PCR Taqman probes used in the present study

| **<u>Probes</u>** | **<u>Catalog No.</u>** |
| --- | --- |
| *Angptl8* (Mouse) | Mm01175863_g1 |
| *B2m* (Mouse) | Mm00437762_m1 |
| *Gapdh* (Mouse) | Mm99999915_g1 |
| *B2m* (Rat) | Rn00560865_m1 |
| *Cebpb* (Rat) | Rn00824635_s1 |

**Table 2.**
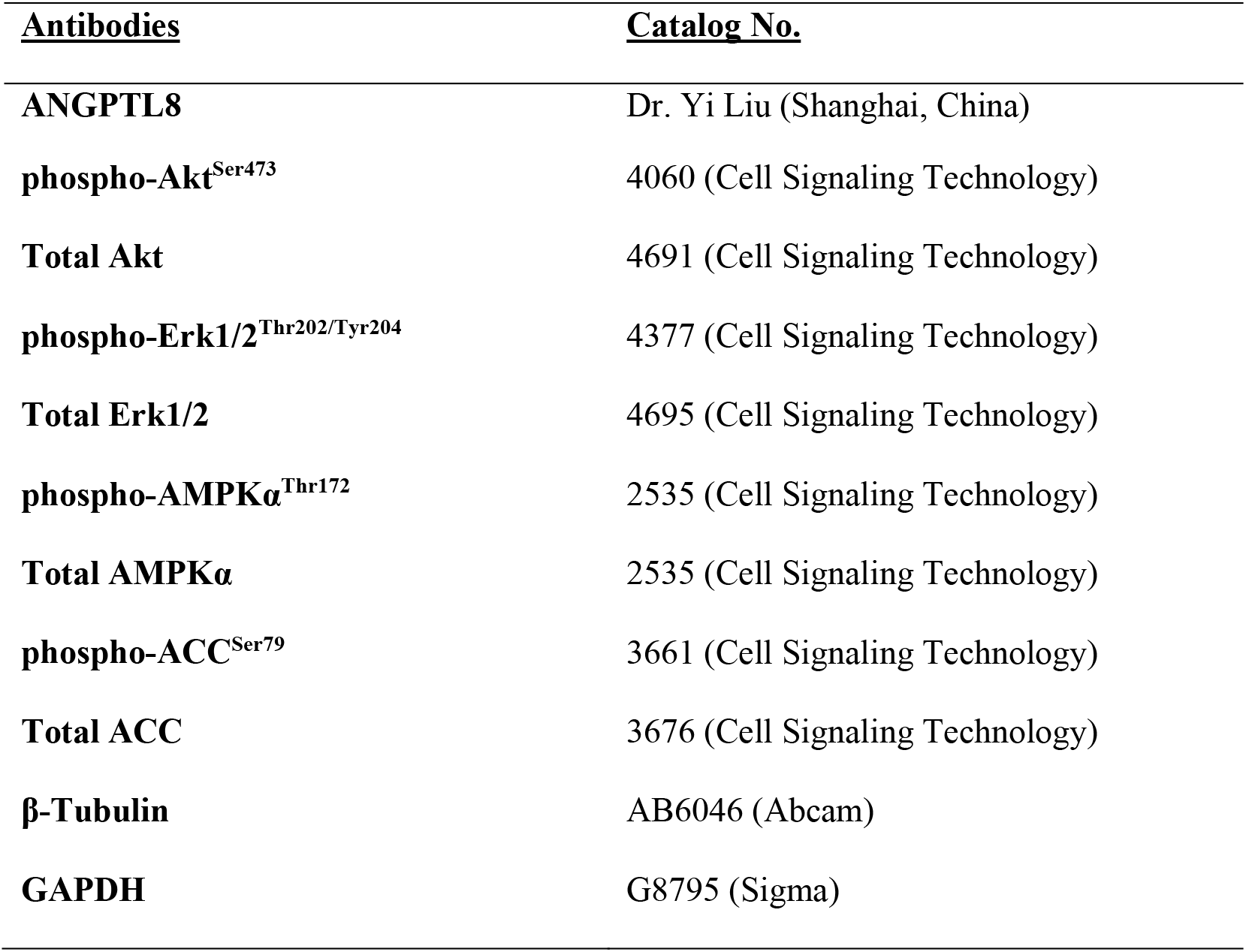
Primer sequences used to generate ANPTL8 promoter constructs

**Table 3.**
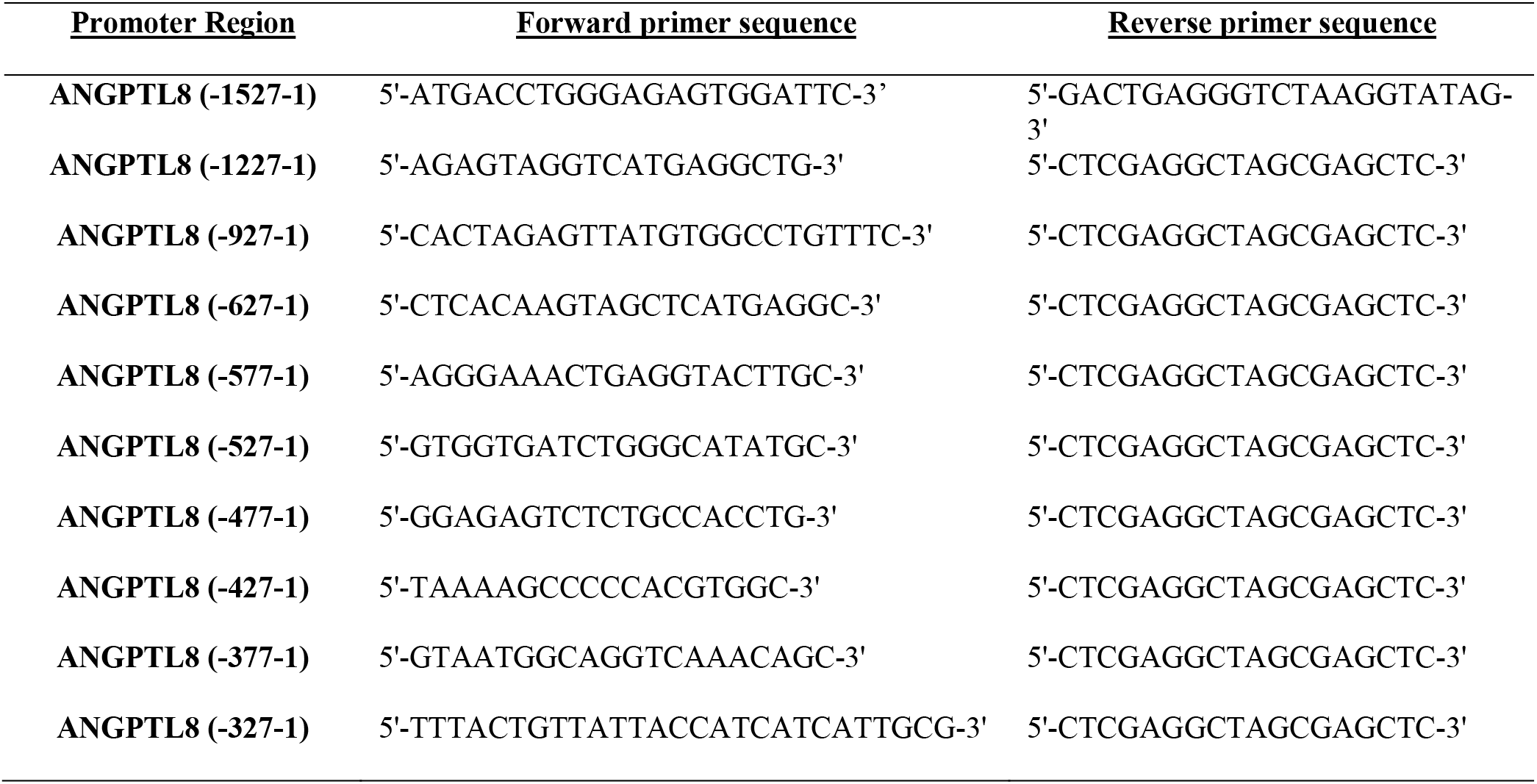
Antibodies used for immunoblotting

### Plasmids Construction and Dual-Luciferase Reporter Assay

The 5’ upstream promotor region of the human ANGPTL8 gene (−1527 to −1, +1 corresponding to start of the first exon) was amplified by PCR and cloned into the pGL4.21-Luc vector (Promega, Madison, WI, USA). Systematic promotor truncations were generated using the Q5 Site-Directed Mutagenesis Kit according to manufacturer’s instructions (New England Biolabs, Ipswich, MA, USA). Plasmid constructs were introduced via Neon electroporation (Invitrogen, Carlsbad, CA, USA) as previously described (16). Following the indicated treatments, cells were harvested for the determination of Firefly and Renilla luciferase activity using the dual-luciferase reporter assay system (Promega), and Firefly luciferase activity was normalized to Renilla luciferase activity. The sequences of all primers are listed in **Table 1**.

### Gene silencing

Short-interfering RNAs (siRNA) against selected genes and scrambled siRNA were purchased from GE Dharmacon (Lafayette, CO, USA). Electroporation of H4IIE cells were performed using Neon Transfection System, as previously described (16).

### Western Blotting and Quantitative Real-Time PCR (qRT-PCR)

Immunoblot analysis was carried out on cell or tissue lysates using primary antibodies described in **Table 2**. Quantitative real-time PCR (qRT-PCR) was performed using pre-designed TaqMan probes (Life Technologies) (**Table 1**). For analysis of ANGPTL8 expression, primers were designed using PrimerQuest Design Tool (Integrated DNA Technologies) and qRT-PCR was performed using SYBR Green. Data were normalized to reference genes *B2m* (for cultured cells and mouse liver) or *GAPDH* (for mouse adipose tissue).

### Statistics

Results are represented as mean ± SEM. Data containing three or more groups was analyzed using one-, two- or three-way anova (adjusted for repeat measures where appropriate) with Holm-Sidak post-hoc testing accounting for multiple comparisons, where appropriate.

## Results

### ANGPTL8 knockout mice

Previous studies have reported that ANGPTL8^−/−^ mice have significantly reduced bodyweight, fat mass and circulating triglycerides (6; 17). In our model we confirmed the effect of ANGPTL8 knockout on fed triglyceride levels, but did not observe any differences in bodyweight or food intake in male (**Fig. 1**) or female (**Supplementary Fig. 1**) mice up to 6 months of age when fed a chow diet. In contrast to some previous studies, we observed an improvement in glucose tolerance in chow fed ANGPTL8^−/−^ male mice (**Fig. 1**) that was associated with a small but significant reduction in fed hepatic triglyceride content (**Supplementary Fig. 3**).

**Figure 1:**
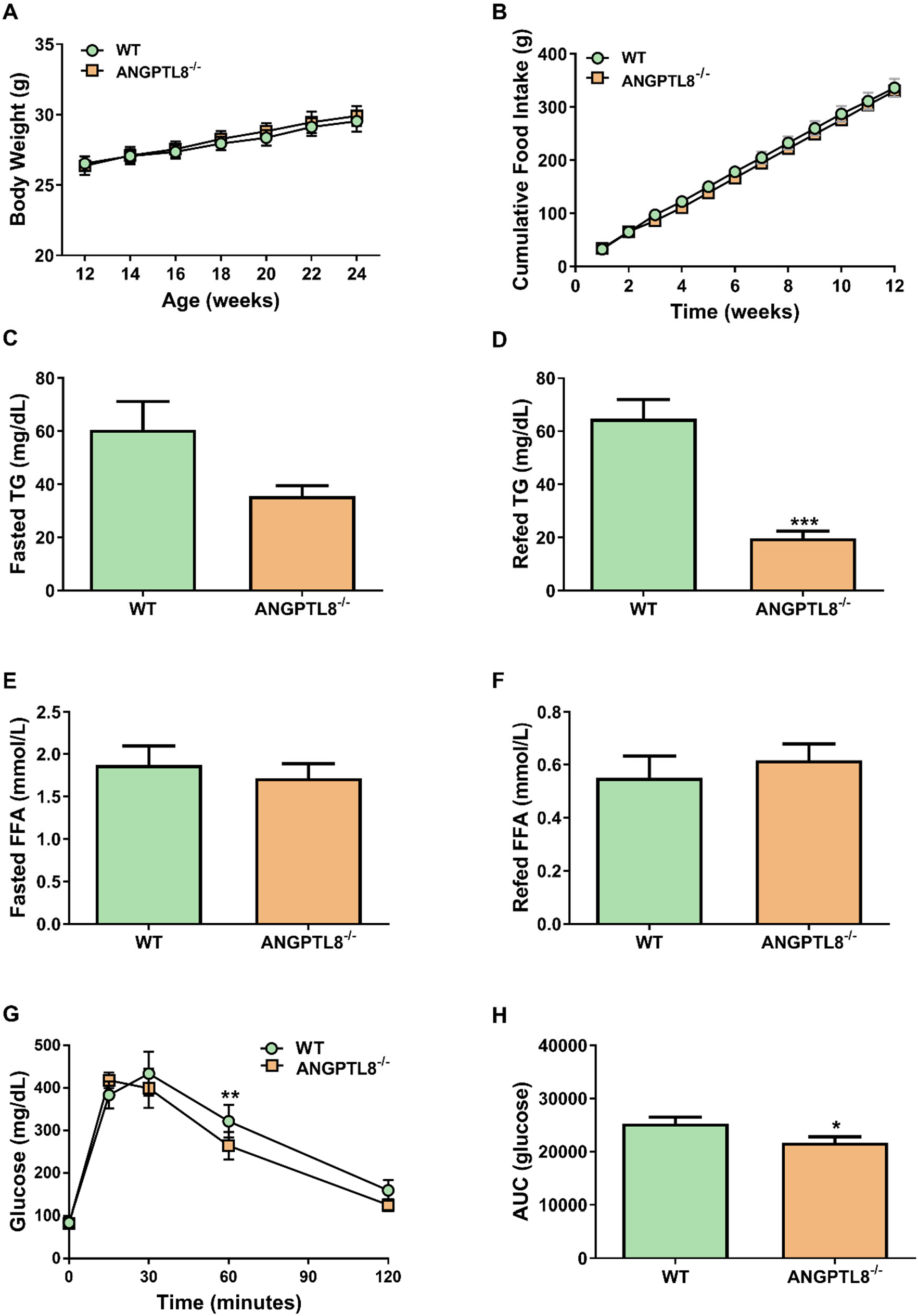
Body weight (A), food intake (B), fasted and refed TG (C and D) and FFA (E and F), glucose tolerance (G) and area under the glucose tolerance curve (AUC; H) were measured in male ANGPTL8 knockout mice (ANGPTL8^−/−^) fed a chow diet for 12-weeks (n = 8 – 17 per group). ^*^p<0.05, ^**^p<0.01, ***p<0.001 vs. WT control mice.

Following a 12-week HFD, both male (**Fig. 2**) and female mice (**Supplementary Fig. 2**) had moderate but significantly improved glucose tolerance without overt changes in weight gain or food intake. Interestingly, this was associated with reduced fed and fasted circulating triglycerides in male (**Fig. 2**) and female mice (**Supplementary Fig. 2**), and lower fed circulating FFAs in male mice only (**Fig. 2**). Interestingly there was no change in hepatic triglycerides following the high fat diet in male mice (**Supplementary Fig. 3**).

**Figure 2:**
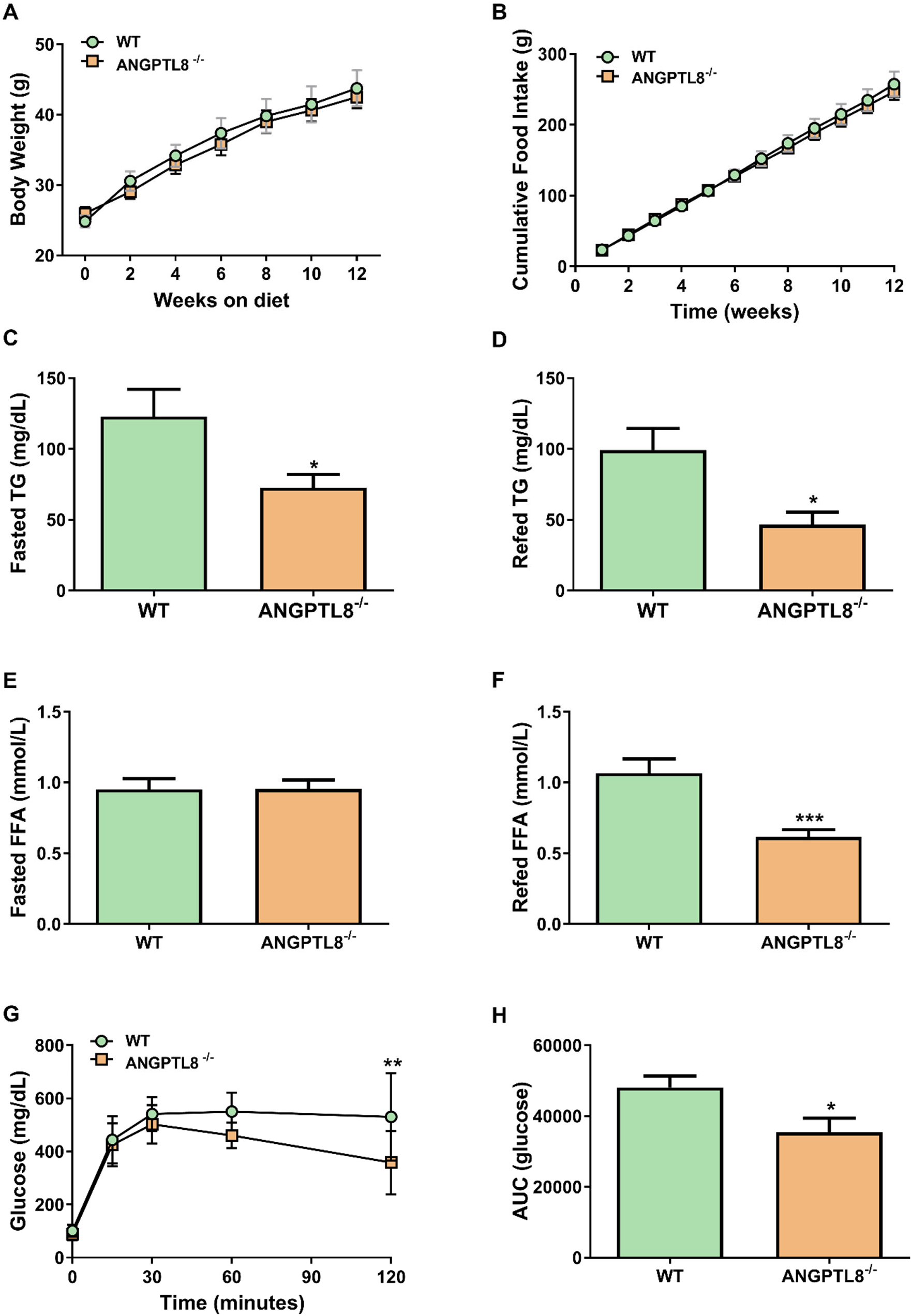
Body weight (A), food intake (B), fasted and refed TG (C and D) and FFA (E and F), glucose tolerance (G) and area under the glucose tolerance curve (AUC; H) were measured in male ANGPTL8 knockout mice (ANGPTL8^−/−^) fed a high-fat diet (HFD) for 12-weeks (n = 7 – 12 per group). ^*^p<0.05, **p<0.01, ^***^p<0.001 vs. WT control mice.

### Regulation of ANGPTL8 expression by insulin and glucose

Given the phenotype of the ANGPTL8^−/−^ mice, we next investigated the physiological and molecular pathways regulating ANGPTL8 expression. Because nutritional status is a powerful regulator of ANGPTL8, we first examined the effect of insulin and/or glucose in cultured hepatocytes and differentiated adipocytes on ANGPTL8 expression. In the rat H4IIE hepatic cell line insulin rapidly upregulated ANGPTL8 mRNA in a dose-dependent manner (**Fig. 3**). Significant upregulation of ANGPTL8 occurred as early as 60 mins following insulin stimulation, and could be detected with an insulin dose as low as 1 nM (**Fig. 3**). Hyperglycemia, alone or in combination with insulin, did not affect ANGPTL8 expression in hepatocytes (**Fig. 3**). In differentiated 3T3-L1 adipocytes, increased ANGPTL8 mRNA was observed with insulin concentrations as low as 100 pM (**Fig. 3**). Consistent with our hepatocyte data, hyperglycemia alone did not increase ANGPL8 expression. However, the combination of hyperglycemia and insulin further increased ANGPTL8 mRNA compared to insulin alone in 3T3-L1 cells (**Fig. 3**).

**Figure 3:**
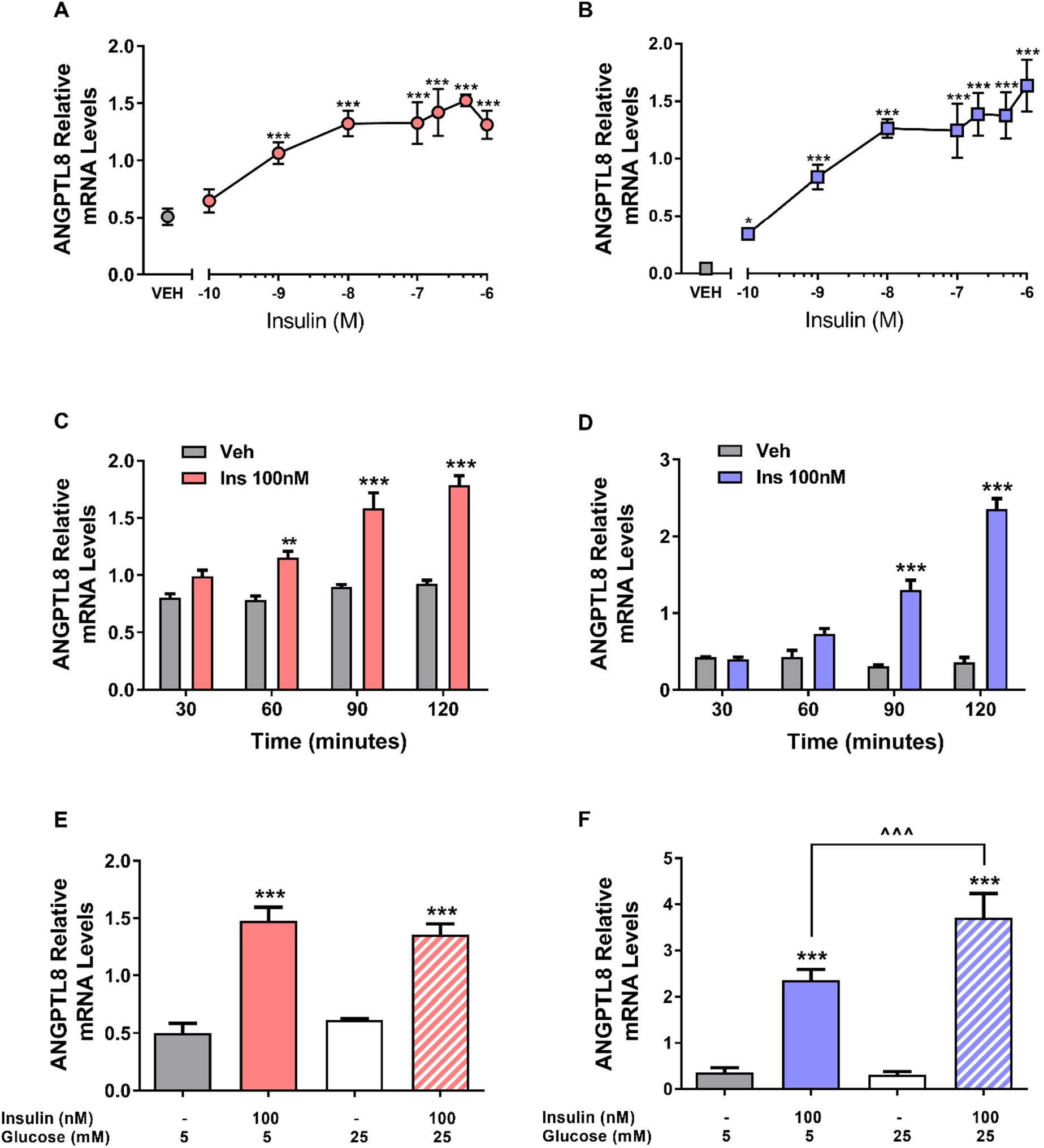
Effect of insulin and glucose on ANGPTL8 mRNA levels in H4IIE hepatocytes and differentiated 3T3-L1 adipocytes. Dose response of ANGPTL8 transcript regulation by insulin in H4IIE (A) and 3T3-L1 (B) cells. Time course of ANGPTL8 transcript regulation by insulin (100 nM) in H4IIE (C) and 3T3-L1 (D) cells. The effect of glucose, alone or in combination with insulin, on ANGPTL8 mRNA levels in H4IIE (E) and 3T3-L1 (F) cells (n = 3 independent experiments). ^**^p<0.01, ^***^p<0.001 vs. VEH or baseline time point; ^^^^^p<0.001 vs. indicated insulin treated cells

To confirm these data in an *in vivo* setting, we employed multiple insulin clamp studies to investigate whether insulin and/or glucose acutely regulate ANGPTL8 levels in the adipose tissue and liver of wild-type mice. In the euglycemic-hyperinsulinemic clamps, ANGPTL8 mRNA was significantly upregulated in liver, but not adipose tissue (**Fig. 4**). In contrast during hyperglycemic-hyperinsulinemic clamps, ANGPTL8 mRNA was increased in both liver and adipose tissue (**Fig. 4**). To isolate the effect of hyperglycemia, we performed hyperglycemic clamps with somatostatin infusion to block endogenous insulin release. In these experiments ANGPTL8 mRNA was not increased in liver or adipose tissue (**Fig 4**). These findings demonstrate that insulin is a key determinant of ANGPTL8 expression in hepatocytes and adipocytes, and suggest that insulin-stimulated glucose uptake plays a unique role in regulating adipocyte ANGPTL8 mRNA.

**Figure 4:**
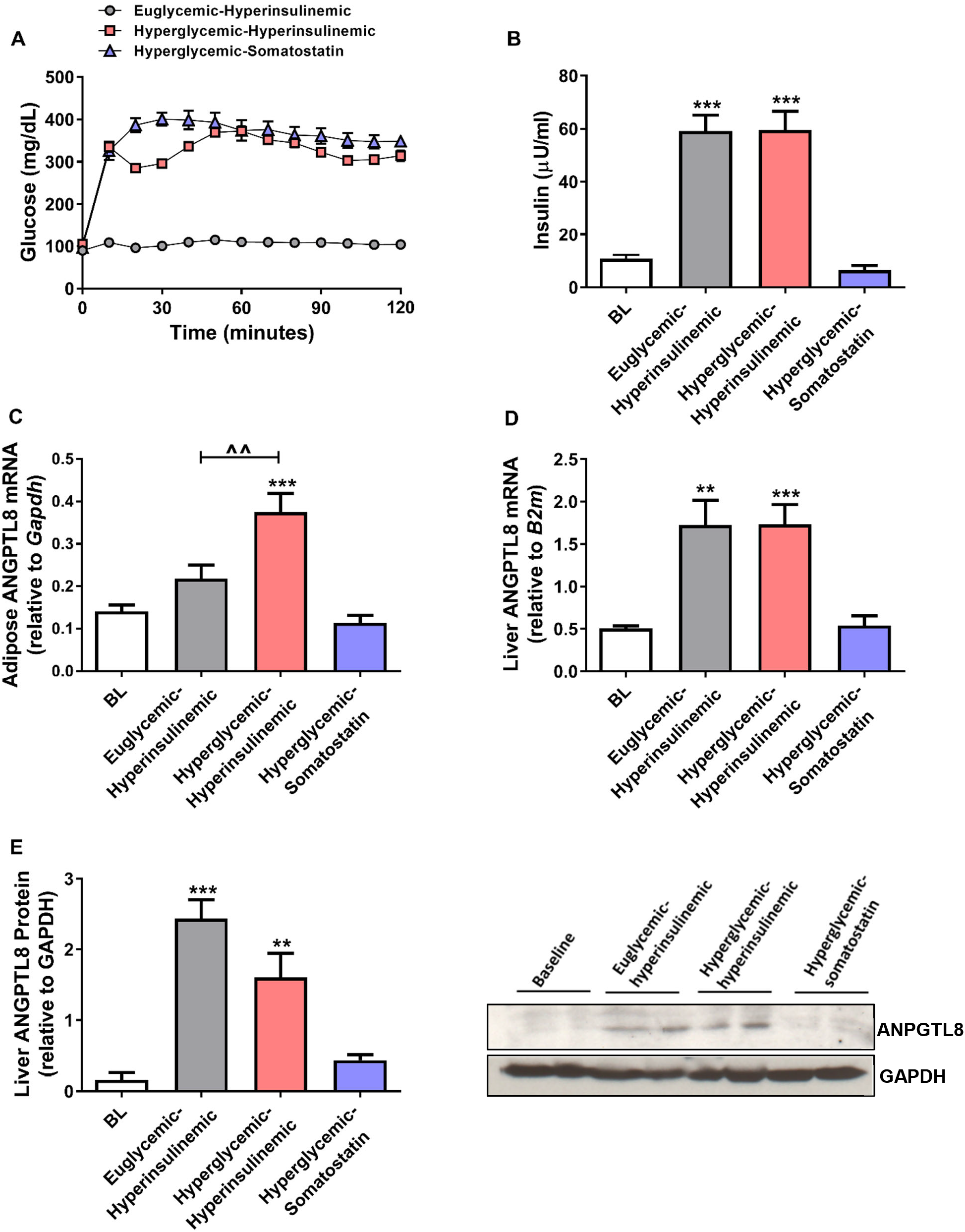
Regulation of ANGPTL8 gene and protein levels by insulin and glucose in mouse liver and adipose tissue evaluated by euglycemic-hyperinsulinemic, hyperglycemic-hyperinsulinemic and hyperglycemic clamps with somatostatin infusion. Plasma glucose and steady-state insulin levels during each clamp are shown (A and B). The mRNA levels of ANGPTL8 in adipose tissue (C) and liver (D) was quantified in tissues collected at the end of each clamp and hepatic ANGPTL8 protein levels (E) were quantified on Western blots (n = 4 – 11/group). ^**^p<0.01, ^***^p<0.001 vs baseline (BL); ^^^^p<0.01 vs. indicated clamp.

### Insulin signaling, AMPK activation and ANGPTL8 expression

We next investigated whether insulin signaling through the canonical PI3K-AKT or the MAPK (ERK) pathways was involved in the regulation of ANGPTL8 mRNA. Surprisingly, neither the PI3K pathway inhibitor wortmannin, nor the MAPK pathway inhibitor PD98059, had any effect on baseline or insulin stimulated ANGPTL8 mRNA expression (**Supplementary Fig 4**). This was despite robust inhibition of both pathways, as demonstrated by significant reductions in protein levels of phospho-AKT^Ser473^ and phospho-p44/42 MAPK^Thr202/Tyr204^ (Erk1/2;) (**Supplementary Fig 4**).

In contrast to the above results, treatment of H4IIE hepatocytes or differentiated 3T3-L1 adipocytes with the AMPK activator 5-Aminoimidazole-4-carboxamide ribonucleotide (AICAR) increased serine 79 (Ser79) phosphorylation of acetyl-CoA carboxylase (ACC) and completely blocked the effect of insulin on ANGPTL8 expression (**Fig 5**). The effect of AMPK activation was more pronounced in hepatocytes, where treatment of cells with AICAR alone modestly buts significantly reduced ANGPTL8 mRNA expression (**Fig 5**)

**Figure 5:**
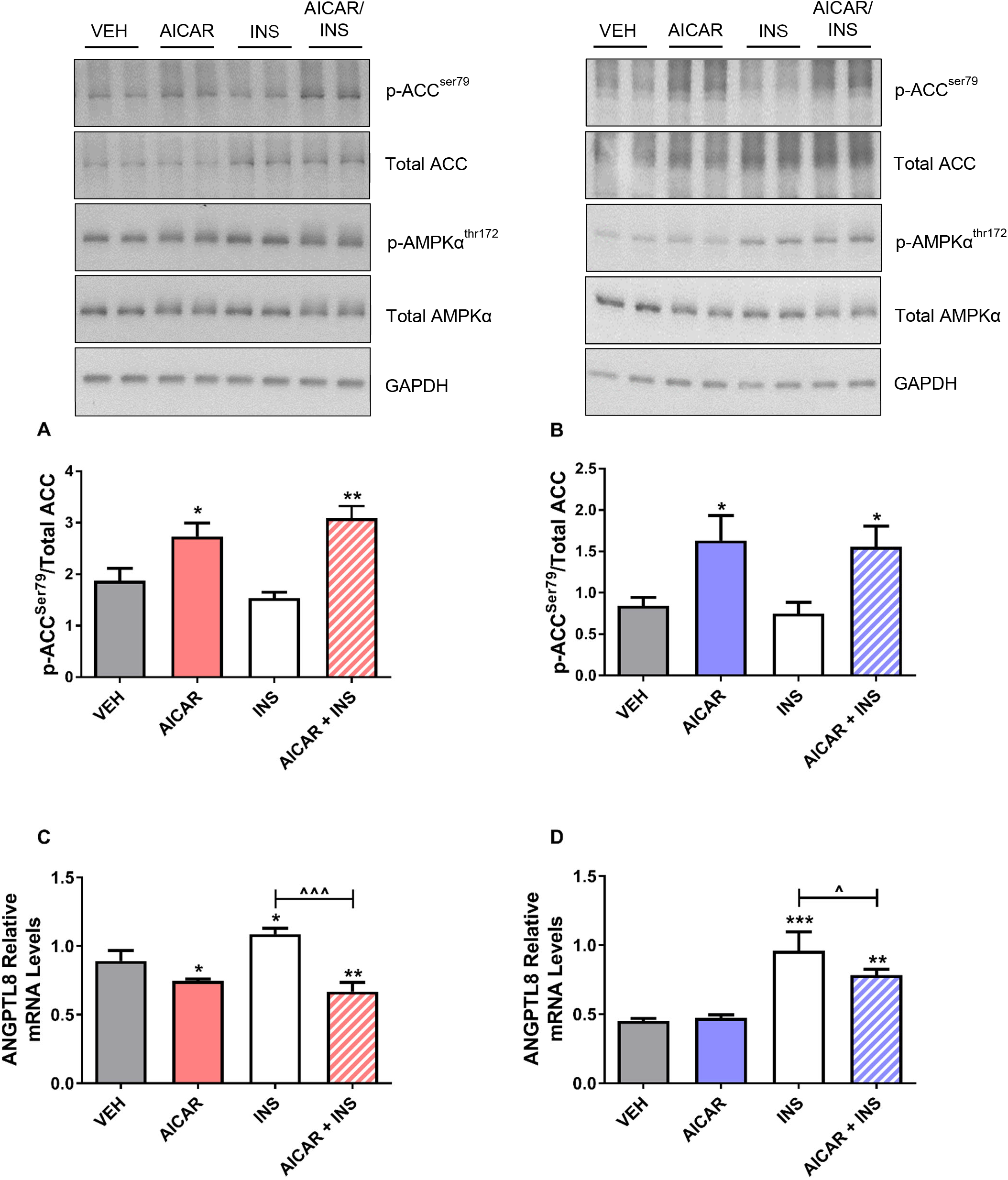
The effect of AMPK activation on ANGPTL8 mRNA levels. H4IIE (A and C) or 3T3-L1 (B and D) cells were incubated with 0.5 mM AICAR in the presence or absence of 100 nM insulin (INS), and ANGPTL8 mRNA levels were evaluated by qRT-PCR. Immunoblotting confirmed the activation of the AMPK pathway (upper panels). Only a single GAPDH loading control is shown for simplicity (n = 3 – 5 independent experiments). ^*^p<0.05, ^**^p<0.01, ^***^p<0.001 vs. VEH control; ^^^p<0.05, ^^^^^p<0.001 vs. insulin only treated cells.

### Transcriptional Regulators of ANGPTL8

Lastly, we focused on the identifying insulin-responsive transcription factors that may regulate ANGPTL8 mRNA expression in hepatocytes. The immediate 1.5 kb promotor region of the human *ANGPTL8* gene was cloned into the pGL4.21-Luc vector and transfected into hepatocytes. Consistent with our earlier observations, insulin treatment significantly increased the activity of the 1.5 kb promoter of ANGPTL8 (**Fig 6**). To isolate specific regions of the promoter that were driving the insulin response, a series of truncations of the promoter were made. The induction of the ANGPTL8 promoter by insulin remained unchanged until a region upstream of −527 bp was removed. Further analysis revealed that this promoter region contained a strong CCAAT/enhancer-binding protein beta (C/EBPβ) consensus binding sight (**Fig 6**). Critically, C/EBPβ expression was significantly increased by insulin treatment in hepatocytes, and in follow-up siRNA studies was found to be required for insulin-mediated ANGPTL8 mRNA upregulation (**Fig 6**).

**Figure 6:**
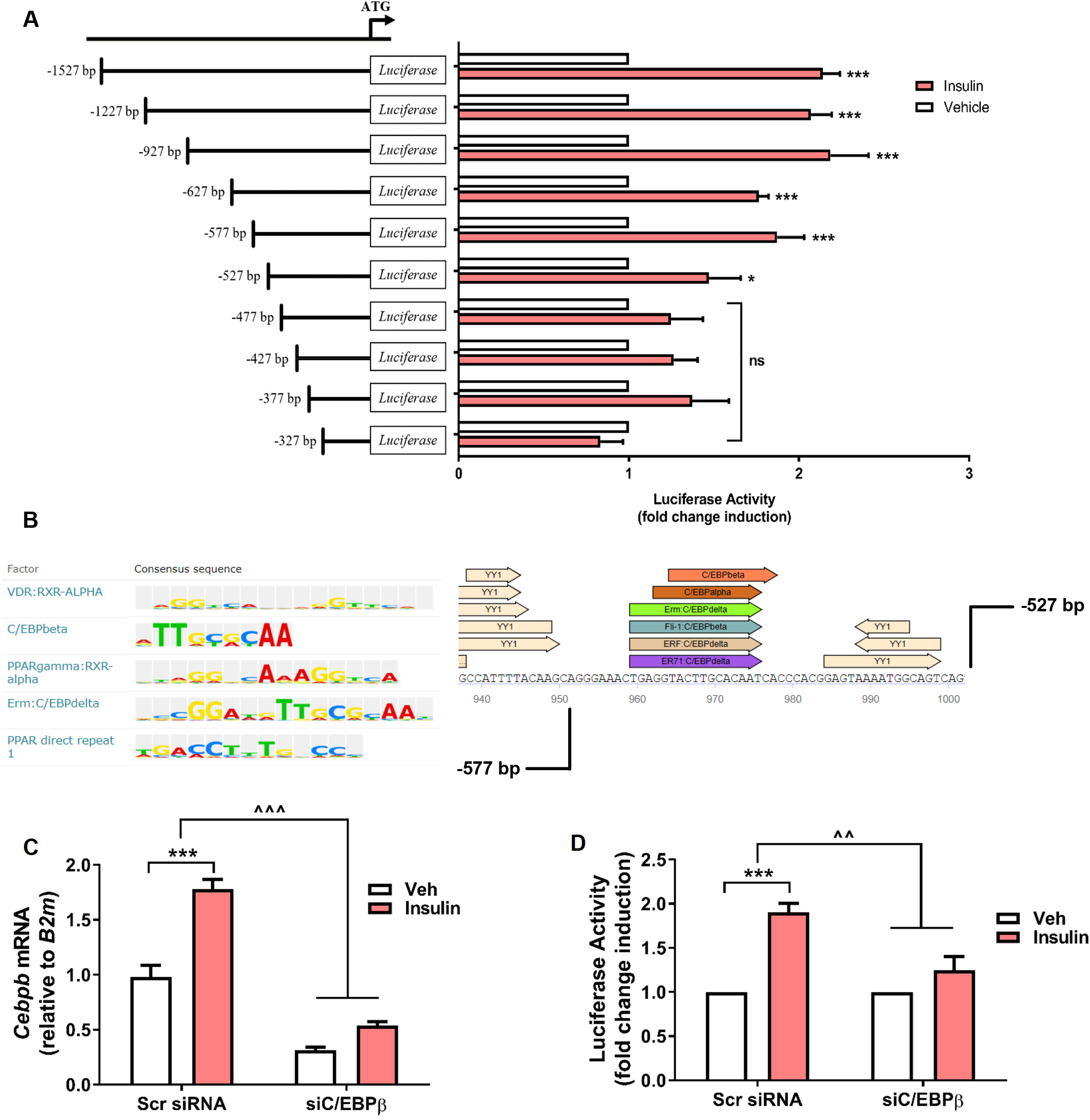
The identification of insulin-responsive transcriptional factor binding sites on the ANGPTL8 promoter using the luciferase reporter system in H4IIE cells. Cells were transfected with pGL-4.2 reporter plasmids containing serial truncated fragments of the ANGPTL8 gene promoter (+1, translation start site) and luciferase activity was detected following incubation with or without insulin (A). Predicted C/EBPβ binding sequence on the ANGPTL8 promoter (B). Cells were co-transfected with pGL-4.21 reporter plasmids containing the 577 bp ANGPTL8 promoter region and scrambled siRNA or siRNA against C/EBPβ and were incubated with insulin for the assessment of C/EBPβ mRNA levels (C) and ANGPTL8 promotor luciferase activity (D). For luciferase assays, the data are corrected to basal luciferase activity of each construct (n = 3 – 5 independent experiments).

## Discussion

In this study we evaluated the metabolic function of ANGPTL8 and its nutritional and hormonal regulation. We first show that ANGPTL8 knockout has a significant effect on glucose tolerance that is independent of weight loss but is associated with reduced circulating triglycerides in mice. We also demonstrate that insulin and glucose are key regulators of ANGPTL8 expression in adipose tissue and liver *in vitro* and *in vivo*, and highlight an important suppressive role for AMPK signaling on ANGPTL8 expression. Taken together these data provide new insights into the regulation and function of ANGPTL8 in two metabolic tissues important in the pathogenesis of obesity and T2DM.

Since the discovery that loss-of-function mutations in the ANGPTL3 gene in humans are associated with combined hypolipidemia (5), there has been considerable interest in targeting the ANGPTL proteins for the treatment of dyslipidemia. However, whether ANGPTL8 represents a viable target for weight loss and/or insulin resistance and T2DM remains controversial (18). In mice ANGPTL8 knockout led to a significant reduction in bodyweight and fat mass in female mice (6) but had no impact on whole-body glucose homeostasis (6; 10). In contrast, we demonstrate that ANGPTL8 knockout had no effect on bodyweight in chow- or HFD-fed male or female mice, but did improve glucose tolerance. This was more pronounced after the HFD, where lower peripheral insulin resistance was observed in male and female mice. Our findings are more in line with a recent study that used antisense oligonucleotides (ASOs) to knockdown ANGPTL8 expression in mice, an intervention that reduced HFD-mediated insulin resistance (11). The explanation for the discrepancy between these rodent studies is not immediately apparent but may reflect the genetic backgrounds of the ANGPTL8 knockout mice. For example, the mice used in the study by Wang et al were generated on a C57BL/6NTac background, whereas the mice used in the present study were generated on a C57BL/6J background. Additionally, it is possible that increased energy expenditure, which we did not measure in the present study, promotes weight loss in some strains of mice lacking ANGPTL8. In support of this hypothesis, ANGPTL8 blockade with a monoclonal antibody increased energy expenditure and reduced weight gain in mice fed a high-fat high-cholesterol diet (19). Despite these differences, further study is required in animal models and humans to definitively determine the impact of ANGPTL8 ablation on both bodyweight and glucose metabolism.

The precise mechanistic role of ANGPTL8 in regulating circulating triglyceride levels remains to be fully elucidated, but it is clear that ANGPTL8 is expressed primarily in adipose tissue and liver and is upregulated in the fed state (14). This contrasts with ANGPTL4, which is reduced in the fed state but increased during fasting (20; 21). The expression of ANGPTL8 also is elevated during adipogenesis *in vitro* and in Ob/Ob mice (14), both of which are hyperinsulinemic states. Our findings indicate that insulin is indeed a primary driver of ANGPTL8 expression in liver and adipose tissue, which is broadly consistent with recent studies (14; 22). A novel aspect of our work, however, is the additive effect of glucose on ANGPTL8 expression in adipose tissue specifically, which may allow for fine tuning of ANGPTL8 expression and local control of triglyceride metabolism in different tissues. Although there is little evidence that ANGPTL8 modifies LPL activity at physiological concentrations alone, it may facilitate the binding of ANGPTL3 and ANGPTL4 to LPL in liver and adipose tissue, respectively (23; 24). It is hypothesized that the binding of ANGPTL8 to ANGPTL4 during the fed state suppresses its function in adipose tissue, thereby promoting LPL activation and lipid storage (24). Conversely, at the liver increased ANGPTL8 forms complexes with ANGPTL3 to inactivate LPL at the site of oxidative tissues (i.e. skeletal muscle) to promote lipid oxidation (23).

In our culture experiments, we observed rapid increases in ANGPTL8 mRNA expression that were not prevented by blocking the PI3K/AKT pathway. This finding is not consistent with recent work demonstrating that the effect of insulin on ANGPTL8 was blocked following the addition of the PI3K inhibitor LY2924002 (25), and may indicate that insulin upregulates ANGPTL8 through novel mechanisms not directly linked to acute AKT phosphorylation. In contrast, we show that AMPK activation potently counteracts the effect of insulin on ANGPTL8 expression, an effect that was more striking in hepatocytes. These data are consistent with the maximal activation of AMPK in fasted liver (26) and its important role in the regulation of whole-body lipid metabolism (26). These data also suggest that insulin and glucose may upregulate ANGPTL8 expression, in part, through the suppression of AMPK signaling. Previous studies have demonstrated that insulin inhibits AMPK through Ser-485 phosphorylation of AMPK α1/α2 (27; 28). In regards to the transcriptional regulation of ANGPTL8, recent studies reported that ANGPTL8 is a target gene of the transcription factors sterol regulatory element binding protein 1 (SREBP1c) and liver X receptor alpha (LXRα) (29). However, using siRNAs targeting SREBP1c or LXRα, we did not observe notable changes in ANGPTL8 mRNA levels in the presence or absence of insulin (data not shown). This led us to explore alternative transcription factors regulating ANGPTL8 in liver. Our experiments identified CCAAT/enhancer-binding protein beta (C/EBPβ) as a key mediator of insulins effect on ANGPTL8 expression. Like ANGPTL8, the expression of C/EBPβ was acutely increased by insulin, supporting a mechanistic link between ANGPTL8 and C/EBPβ. The observed regulation of C/EBPβ by insulin, and it’s role in regulating ANGPTL8, is consistent with previous reports (30), and with data from C/EBPβ knockout mice, which display impaired adipose tissue development and gross abnormalities in lipid and glucose homeostasis (31; 32). Further studies will be required to interrogate the crosstalk between the insulin signaling pathway and AMPK activation, and to determine the tissue-specific transcriptional pathways that are responsible for the regulation of ANGPTL8 levels.

The finding that insulin with or without hyperglycemia regulates ANGPTL8 expression adds important context to studies that have examined circulating ANGPTL8 levels in humans with metabolic disease, often with conflicting findings. For example, several studies have demonstrated increased ANPGTL8 in patients with obesity (33; 34), whereas others have shown that ANGPTL8 is reduced in obese humans (12). Similarly, in human subjects with T2DM the levels of ANGPTL8 have been shown to be increased (34‒36), decreased (12), or unchanged (37). The reason for these discordant findings is unclear but has been postulated to be the result of suboptimal antibodies and ELISA kits used to measure plasma ANGPTL8 (13; 38). Indeed, attempts by us to measure circulating ANGPTL8 in human plasma by Western blot or ELISA were unsuccessful and neither method met our internal authentication standards. However, a biological explanation is also plausible as it is clear from the current findings that ANGPTL8 levels are highly sensitive to alterations is circulating insulin and glucose concentrations. Consistent with this hypothesis ANGPTL8 levels were positively correlated with 2 hour OGTT insulin and glucose levels in human subjects (39). Interestingly, and in contrast to the results from our clamp experiments, circulating ANGPTL8 levels were reduced in humans during a hyperinsulinemic-euglycemic clamp (40). These data might suggest that ANGPTL8 is indirectly regulated by other circulating factors (i.e. FFA) or, more likely, that ANGPTL8 is not readily secreted from adipose tissue or liver into the plasma. In fact, available evidence suggests that ANGPTL8 is not secreted from tissues unless it is in complex with ANGPTL3 (23; 40) or ANGPTL4 (24). Thus, quantifying ANGPTL8 in human plasma may be of limited value in the absence of concomitant ANGPTL3 and ANGPTL4 quantification.

In summary, our study demonstrates that ANGPTL8 plays an important metabolic role in mice that may extend beyond triglyceride metabolism and that ANGPTL8 may regulate hepatic and peripheral insulin sensitivity and glucose homeostasis. We have also shown that insulin and glucose play key roles in regulating tissue-specific ANGPTL8 expression and reveal that AMPK activation is an important negative regulator of ANGPTL8 in adipose tissue and liver.

## Acknowledgements

We thank Dr. Ren Zhang at Wayne State University for providing the ANGPTL8^−/−^ mice and Dr. Li Yiu at the Chinese Academy of Sciences for providing the ANGPTL8 antibody.

## Author Contributions

The study was designed by L.N and LZ. and the data analysis, conclusions and manuscript drafting were carried out by L.N and LZ. C.E.S., T.M.B, M.A.G and M.F. contributed to the data generation and approved the final manuscript. All authors contributed to the discussion of the results.

## Supplementary Figure Legends

**Supplementary Figure 1:**
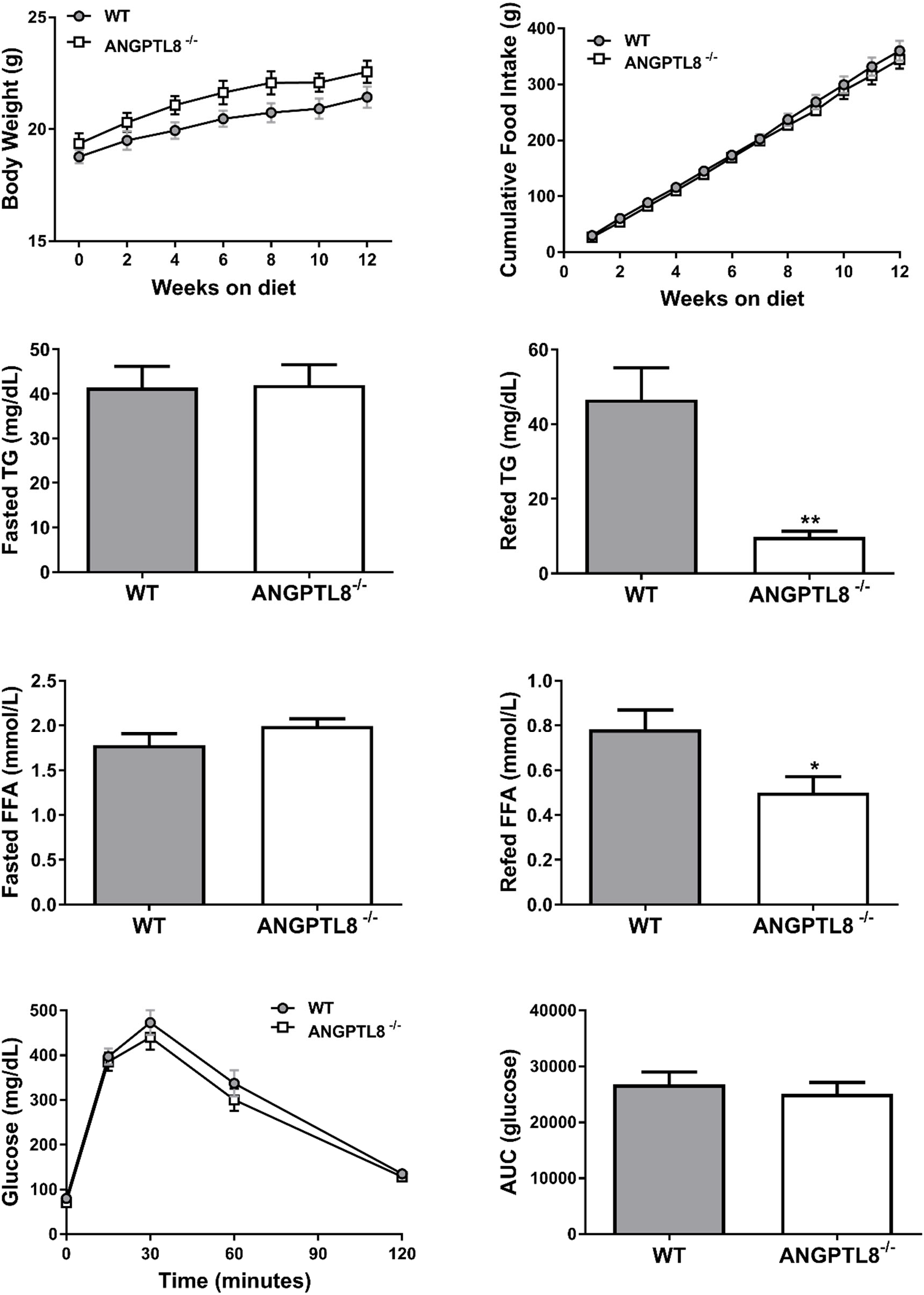
Body weight (A), food intake (B), fasted and refed TG (C and D) and FFA (E and F), glucose tolerance (G) and area under the glucose tolerance curve (AUC; H) were measured in female ANGPTL8 knockout mice (ANGPTL8^−/−^) fed a chow diet for 12-weeks (n = 8 – 12 per group). *p<0.05, **p<0.01, ***p<0.001.

**Supplementary Figure 2:**
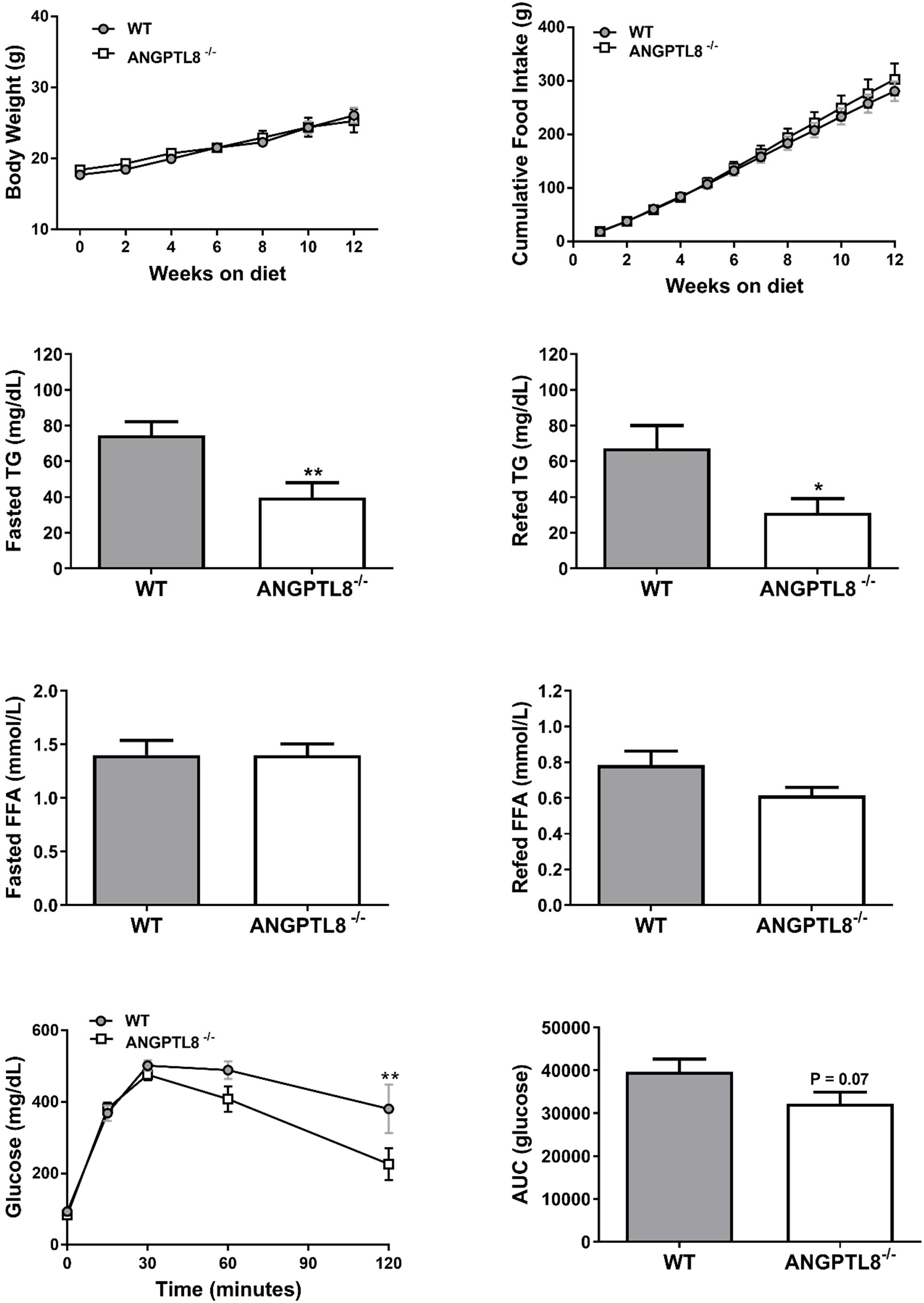
Body weight (A), food intake (B), fasted and refed TG (C and D) and FFA (E and F), glucose tolerance (G) and area under the glucose tolerance curve (AUC; H) were measured in female ANGPTL8 knockout mice (ANGPTL8^−/−^) fed a high-fat diet (HFD) for 12-weeks (n = 5 – 14 per group). *p<0.05, **p<0.01, ***p<0.001.

**Supplementary Figure 3:**
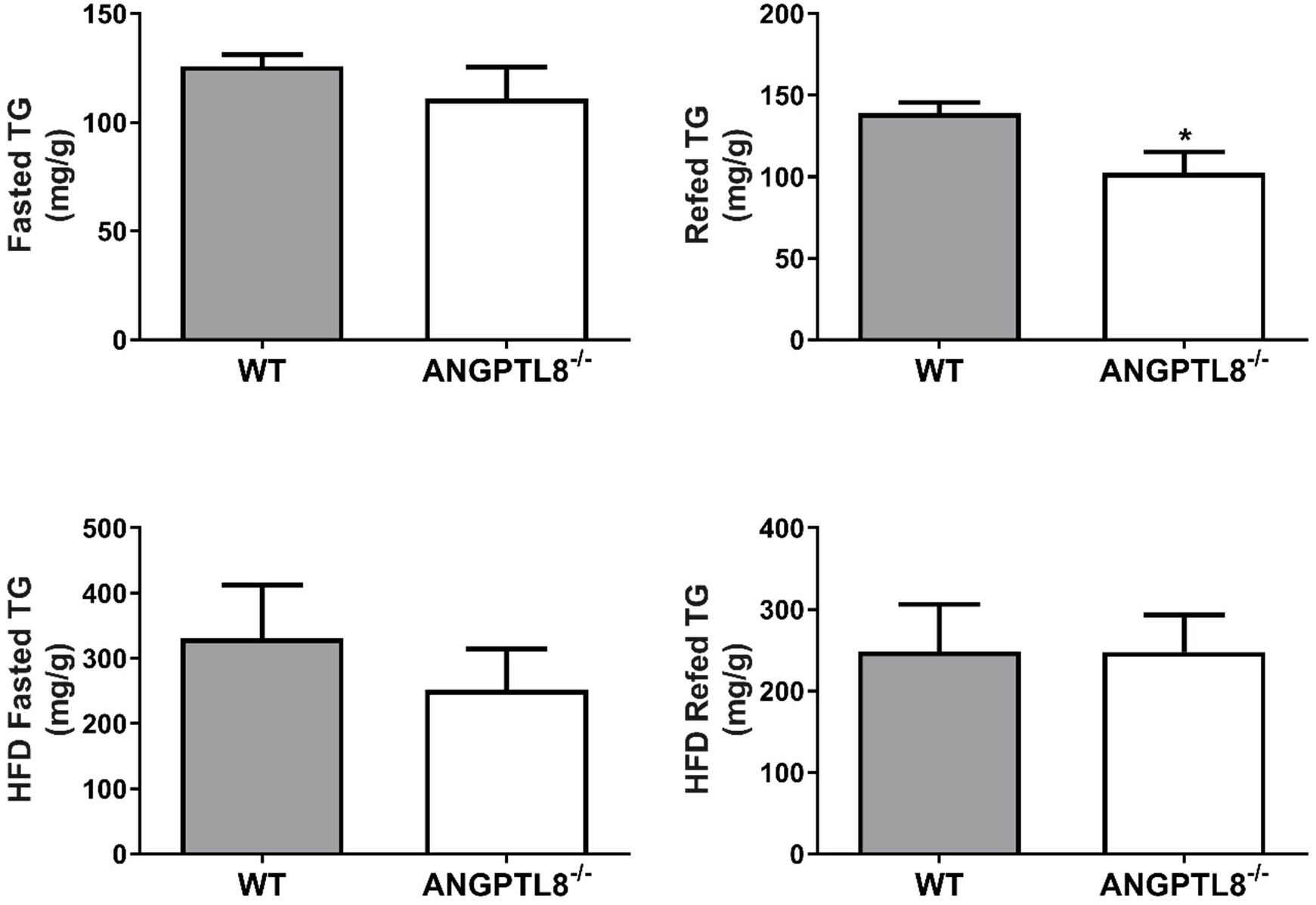
Fasting and refed heaptic triglycerides were quantified in male ANGPTL8-/- mice following chow or high fat feeding (n = 4 – 5/group). *p<0.05 vs. control

**Supplementary Figure 4:**
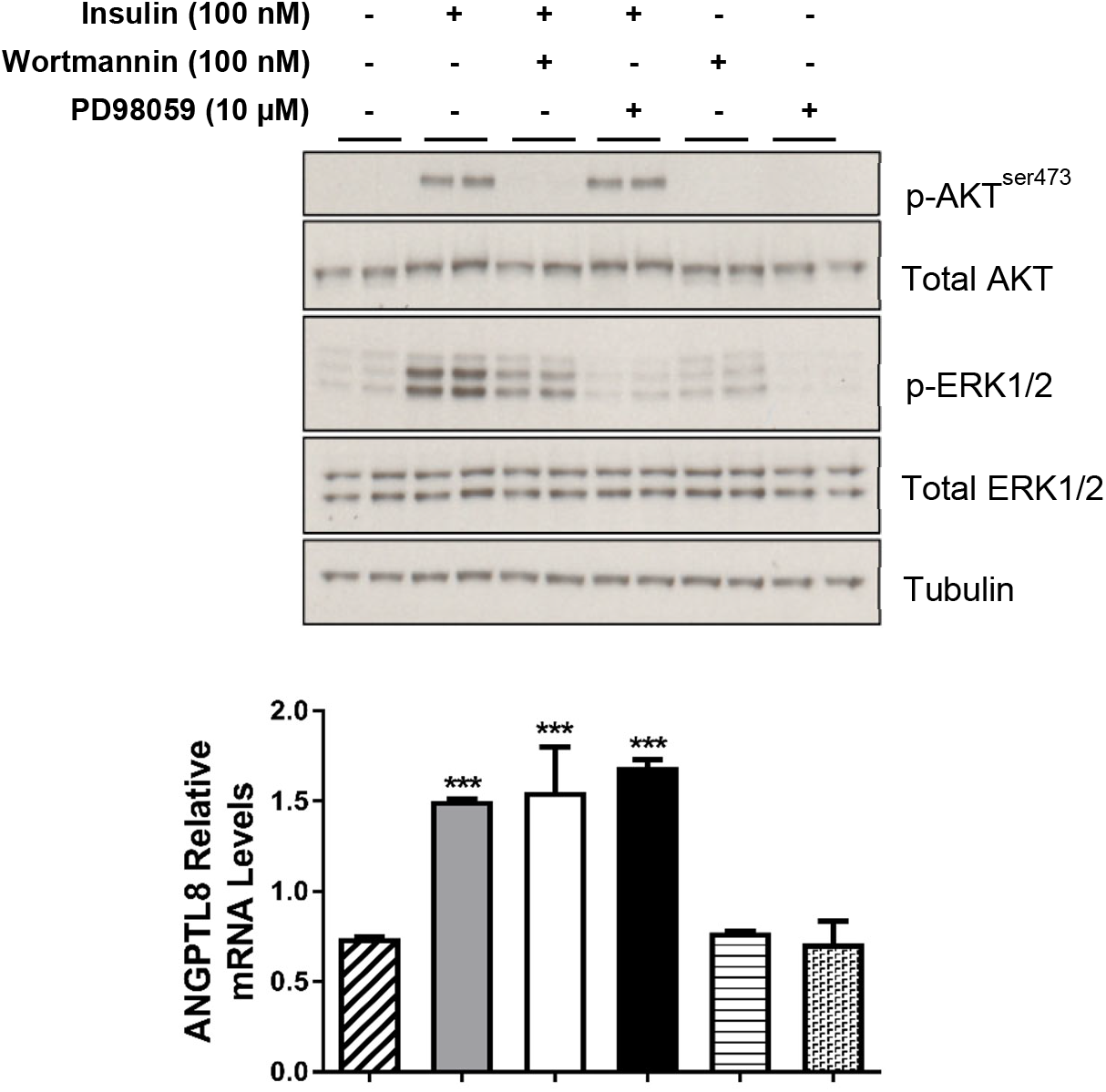
Effects of wortmannin and PD98059 on ANGPTL8 gene expression in the presence of insulin in H4IIE cells. H4IIE cells were incubated with 100 nM wortmannin or 10 uM PD98059 in the presence of 100 nM insulin and ANGPTL8 mRNA levels were evaluated by qRT-PCR. Immunoblotting confirms the inhibitory effects of wortmannin and PD98059 on the PI3K and MAPK pathways, respectively (n = 3 independent experiments). ***p<0.001 vs. VEH control

